# Integrating microfluidic automation into thermoplastic devices for analysis of small volumes of blood

**DOI:** 10.1101/2025.06.28.662160

**Authors:** Josue U. Amador-Hernandez, Alan M. Gonzalez-Suarez, Gulnaz Stybayeva, Gabriel A. Caballero-Robledo, Jose L. Garcia-Cordero, Alexander Revzin

## Abstract

Microfluidic devices hold considerable promise for miniaturizing and automating laboratory scale workflows involving sample preparation and biomarker analysis. By-and-large microfluidic automation has been implemented using polydimethyl siloxane (PDMS) – a material that presents challenges for scalable manufacturing. In this paper, we fabricated an automated microfluidic device composed of thermoplastic material, poly (methyl methacrylate) (PMMA), and demonstrated a sophisticated workflow consisting of plasma extraction, sample aliquoting and sample-reagent mixing steps performed in the device. Automation was accomplished using multiple on-board microvalves consisting of a pressure sensitive adhesive (PSA) membrane sandwiched between PMMA flow and control layers. In addition to novel microvalves, our device included features designed to address challenges arising from the gas impermeable nature of PMMA. These features included a bubble trap and air micro-tanks to supply O_2_ for enzymatic reactions. We rigorously characterized the performance of the microfluidic device and demonstrated that the quality of plasma extracted in the device is comparable to a standard plasma isolation method involving centrifugation. We also showed close agreement between glucose levels measured in the device and by standard assay. As a final demonstration of utility, we loaded 5 µL of blood, isolated plasma and detected three analytes simultaneously - glucose, lactate and H_2_O_2_. In addition to minimizing the volume of blood and eliminating standard laboratory equipment (e.g., centrifuge), our device enabled a rapid response (<10 min). The use of thermoplastic materials may pave the way for inexpensive, mass-produced automated microfluidic devices for rapid detection of blood biomarkers at the point of care.

## INTRODUCTION

There is a growing interest in collecting information about patient health to make better informed decisions about disease diagnosis and treatment. Microfluidic devices may allow for rapid and multiplexed analysis based on a small volume of a biological fluid (e.g., blood) and may be applied to multiple diseases and medical conditions. Neonatology is one medical area that stands to benefit from multiplexed analysis of blood or other biological fluids. Newborns, particularly premature babies, are at risk of hypoglycemia, infections and other health complications.^1^ While the total blood volume for a premature baby may be as small as 50 mL^2–5^, the laboratory assays in use today are not designed to be volume-sparing and require collecting >500 µL of blood or up to 1% of the total blood volume.^6,7^ Given the frequent blood draws required during a health crisis, newborns are at high risk of anemia.^6–8^ There is therefore a strong need to develop sample-sparing analytical approaches for neonatology applications. While there are commercial glucometers designed to work with a finger prick of whole blood, they are unreliable in the hypoglycemic range experienced by infants (< 2 mM).^5,9,10^ Because of this, neonatal glucose levels are typically measured in plasma isolated using standard laboratory equipment in a laborious process that takes 30 to 60 min to complete and requires ∼500 µL of blood.^6,7^ An automated microfluidic device capable of isolating plasma and detecting glucose (as well as other biomarkers) rapidly and at the point of care is therefore beneficial.

An early example of microfluidic automation was work by Quake and co-workers who integrated pressure-actuated microvalves into PDMS microfluidic devices in the early 2000s.^11^ It is well-established that such on-board microvalves (as opposed to external valves or pumps) are essential for applications where metering and routing microliter volumes is important.^12^ The vast majority of microfluidic devices with on-board valving capabilities are fabricated using polydimethylsiloxane (PDMS). This material has multiple advantages. It is gas permeable, biocompatible and elastomeric, making it well-suited for fabricating deformable structures/membranes that serve as microvalves. However, PDMS is a thermosetting material which poses challenges to scalable manufacturing. There has been an increasing interest in fabricating microfluidic devices using thermoplastics, such as cyclic olefin copolymer (COC), poly(methyl methacrylate) (PMMA), and polyethylene terephthalate (PET).^13–16^ These materials allow large-scale and low-cost production using techniques such as injection or compression molding, microthermoforming by rolling, casting, micromilling, and laser ablation.^17–21^ However, the integration of flow control systems (valves and pumps) into thermoplastic devices has historically been a challenge because the rigidity of thermoplastics makes it difficult to create flexible, deformable membranes necessary for effective valve operation.^22^ In some cases, thermoplastic valves incorporated such materials as PDMS to provide deformability.^23^ Our team recently described a strategy of using pressure sensitive adhesive (PSA) for multi-layer bonding of PMMA.^24^ In this strategy, PSA was sandwiched between the PMMA control and flow layers, serving as both an adhesive layer and an actuator membrane of on-board microvalves.

In this paper, our goal was to fabricate a microfluidic device using low-cost thermoplastic materials such as PMMA for automated extraction of plasma and analysis of multiple biomarkers while using 5 µL of blood and providing an answer within 10 min. This approach offers a cost-effective, rapid, and reliable solution for point-of-care diagnostics, with potential for large-scale production and widespread accessibility.

## MATERIALS AND METHODS

### Fabrication of the automated microfluidic device

1.6-mm thick PMMA sheets (8560K179, McMaster-Carr) were used to produce the microfluidic device. The PMMA sheets were carved using a milling machine (MDX-50, Roland AG) with a spindle speed of 13,500 rpm and a feed rate of 0.1 mm/s. A 127 µm square end-mill (SGS02275, Kyocera) was used to carve the microchannels and valve structures in the flow layer, while a 200 µm square end mill (SGS02204, Kyocera) was used to carve out the control lines in the control layer. An 800 µm square end mill (8915A486, McMaster-Carr) was used to drill holes as inlets/outlets and cut individual microfluidic devices from the PMMA sheet. The flow and control layers were aligned and bonded using double-sided tape (pressure sensitive adhesive, PSA, ARcare 92712, Adhesive Research). The assembly process and gate valve passivation were carried out as described in our previous work.^24^ Briefly, one of the protective liners of the PSA is peeled off and the exposed adhesive layer bonded to the control layer. Next, the second protective liner is removed and a drop of clear nail varnish diluted in acetone at a 1:10 ratio is placed in regions overlapping the area of the gate valve. Afterwards, the flow layer is aligned and bonded to the control layer by applying pressure.^24^

### Operation of the automated microfluidic device

The device was integrated with thirteen normally closed gate valves that were actuated with six pneumatic lines to carry out experimental workflow involving plasma separation, sample aliquoting and mixing. The six pneumatic control lines were connected by Tygon tubing (outer diameter 1.5 mm, AAD04103, Saint Gobain) to custom-made pneumatic controller. The system uses an Arduino microcontroller (Arduino Elegoo Mega 2560) to manage a rack of sixteen relays (SainSmart 16-Channel Relay Module, Amazon, B01N6JFQZW), which in turn is connected to electropneumatic solenoid valves (MH1 miniature valve, Festo, Germany). The three-way external solenoid valves are connected to house air, a vacuum line and the device. A manual regulator (11142NNKRSP, Fairchild) was used to apply 138 kPa to close the gate valve and −95 kPa to open it. The positive pressure applied to the control layer of the microfluidic device caused the PSA membrane to deform downward and close the gate valve. Conversely, applying negative pressure caused the membrane to deform upward and open the gate valve. The microfluidic device was integrated with thirteen micro valves, operated through six pneumatic lines connected to the solenoid valves of the PNC by Tygon tubes. The three-way solenoid valves were configured as follows: port 1 was connected to the pressure source (air compressor), port 2, which controls the actuator, was connected to the control lines of the microfluidic device, while port 3 (exhaust) was linked to a vacuum pump. This allowed the opening of the PMMA valves and facilitates the escape of air from port 1 when the solenoid valve is de-energized.

A custom LabVIEW interface was used to send commands (via serial) to a custom C++ algorithm running on the Arduino board to control and automate functions which includes a scheduling function that allows users to set predefined schedules for executing specific commands on the microfluidic device, such as opening and closing gate valves, active mixing, or pumping reagents at different frequencies ranging from 3.3 to 20 Hz.

### Characterization of the plasma separation module

The plasma separation module of the automated device contained a sample port for dispensing whole blood, a channel for plasma collection and an outlet through which negative pressure was applied using a pressure regulator (LU-FEZ-N345, Fluigent, France). The sample port incorporated a plasma separation membrane (PSM) (Cobetter Filtration, Hangzhou, China) for filtering blood cells and extracting plasma from whole blood. The membrane was cut into a circular piece of 4.5 mm in diameter which was sandwiched between control and flow layers. The PSM and the PMMA layers were bonded using double-sided tape (PSA). The whole blood samples (from healthy adults) were obtained in 6 mL ethylenediaminetetraacetic acid (EDTA)-coated tubes from the Mayo Clinic Blood Bank from de-identified donors. The use of these samples did not require the approval of the Institutional Review Board (IRB). Five microliters of whole blood were pipetted onto the sample port and a pressure of −5.51 kPa was applied to the outlet of the device for plasma extraction. The blood cells were retained in the PSM, allowing the extraction of ∼1 µL of plasma. We wanted to characterize the quality of plasma extracted in the microfluidic device using spectrophotometry. This required a volume of ∼20 µL – much larger than what was available in the automated device. We therefore developed a simple microfluidic plasma extraction device, without automation and bioanalysis capabilities, which used similar plasma extraction conditions (negative pressure, plasma separation membrane) but allowed loading 100 µL of blood to extract 20 to 25 µL of plasma. This device was fabricated and operated in a manner similar to the automated microfluidic device described above.

In parallel with the on-chip blood processing, plasma was separated from the same blood sample using a centrifugation-based protocol. Briefly, blood (3 mL) was centrifuged at 1500 ×g for 12 minutes at 5 °C. The blood cells formed a pellet at the bottom of the tube and plasma was pipetted to a vial for storing. The levels of hemoglobin and bilirubin in plasma samples isolated using either a microfluidic device or centrifugation were determined with commercial kits (hemoglobin assay kit MAK115 from MilliporeSigma; and total bilirubin assay kit, BILT3, from Roche). A NanoDrop spectrophotometer (Thermo-Fisher Scientific) was used to make absorbance measurements.

### Detection of analytes in the automated microfluidic device

The bioanalysis module consisted of three analysis units. Each analysis unit contained two 20 nL chambers and was used to mix an aliquot of sample and reagent in a 1:1 ratio. After separation, the plasma was introduced into the sample chambers at the same time with assay components injected into the reagent chambers. The sample and reagent aliquots were homogenized by active mixing with two gate valves operating at 10 Hz for 2 min. This ensured that the enzymatic reaction was not limited by mass transport. Time-lapse micrographs were taken every 10 seconds for 30 min for each assay.

To generate a calibration curve for glucose detection, we prepared a series of glucose solutions (0 to 10 mM) in 1x PBS which were infused into the device. The reagent chambers contained a mixture of glucose oxidase (GOx, G2133-50KU, Sigma Aldrich) at 70 U/mL, horse radish peroxidase (HRP, ICN19537325, Fisher Scientific) at 117 U/mL, ADOS (OC01, Dojindo laboratories) at 3.6 mM and 4-AAP (A4382-100G, Sigma Aldrich) at 3.1 mM. The combining of these reagents with the glucose solution at a 1:1 ratio resulted in a magenta-colored product. The color intensity was measured in arbitrary units (a.u) and was correlated to glucose concentrations. The colorimetric reactions were quantified using an inverted microscope with a color camera (IX83, Olympus, Japan) and images were analyzed using Image J.

Each blood sample was split into two parts, one was used for plasma separation and glucose detection in the microfluidic device, another for analysis using standard approaches. A commercial glucose detection kit (A22189, Thermo Fisher) was used to measure glucose plasma samples. The glucose assay was performed according to the manufacturer’s specifications in a 96-well plate and a plate reader (Synergy H3, BioTek).

The reagents for lactate detection involved enzymatic breakdown by lactate oxidase (LOx) at 4.8 µM (71718-10G, Sigma Aldrich) HRP at 117 U/ml, ADOS at 3.6 mM and 4-AAP at 3.1 mM. Incubation of these reagents with lactate produced a magenta-colored product. Sodium L-Lactate (71718-10G, Sigma Aldrich) was used to construct calibration curves for a dynamic range from 0 to 20 mM. The colorimetric readout was quantified in the manner identical to glucose detection described above. The detection of H_2_O_2_ was carried out using the same reagents included in the glucose detection kit mentioned in the glucose detection section. For the detection of H_2_O_2_, the concentrations of HRP and Amplex Red were adjusted to concentrations of 0.2 U/ml and 5.26 µM, respectively. A calibration curve of H_2_O_2_ was constructed with a dynamic range from 0 to 100 µM. Fluorescence was measured at 590 nm using a fluorescence microscope (IX83, Olympus, Japan).

### Characterization of active mixing

Active mixing was achieved by the sequential activation/deactivation (10 Hz) of two microvalves, one residing on top of the sample chamber, another on top of the reagents chamber, creating chaotic flows in both chambers. The mixing process was assessed using food dye and enzymatic assay for the glucose detection described above. A magenta-colored solution as a product of the enzymatic reactions was perfused into the sample chamber, while 1X PBS was used to fill the reagent chamber. Both solutions were mixed, and the intensity of the magenta color was monitored in both chambers by taking images every 10 seconds for 15 minutes. Control experiments were conducted where the magenta solution and PBS were allowed to mix by diffusion only while taking images every 30 minutes for 12 hours. Mixing was considered complete when the intensity of the magenta color was the same in both the sample and reagent chambers.

### Data analysis

To determine the glucose concentration or evaluate active mixing, the magenta color intensity of the chambers was quantified for each acquired image. A rectangular region of interest of 0.41 mm^2^ in the center of each sample and reagent microchambers was used to measure intensity of the magenta color. The intensity values of the magenta color were obtained by performing a conversion of the RGB image to the CMYK color space (cyan, magenta, yellow and black).

## RESULTS AND DISCUSSION

The goal of this study was to fabricate a thermoplastic microfluidic device to carry out a sophisticated workflow involving plasma separation as well as sample aliquoting and mixing. The device (shown in **Figure 1**) consisted of a plasma separation module and three bioanalysis units (chamber volume of 20 nL) integrated with on-board gate valves. The device received 5 µL of blood, isolated 1 µL of plasma and delivered it into bioanalysis units. Gate valves were used to sequester each bioanalysis unit from its neighbor and to actively mix the contents of sample and reagent chambers within each unit. As described below, we characterized the performance of the device and developed solutions to challenges arising from the use of gas-impermeable thermoplastic material for device construction. The automation and the whole workflow with coloring dyes is highlighted in **Video 1**.

**Figure 1.**
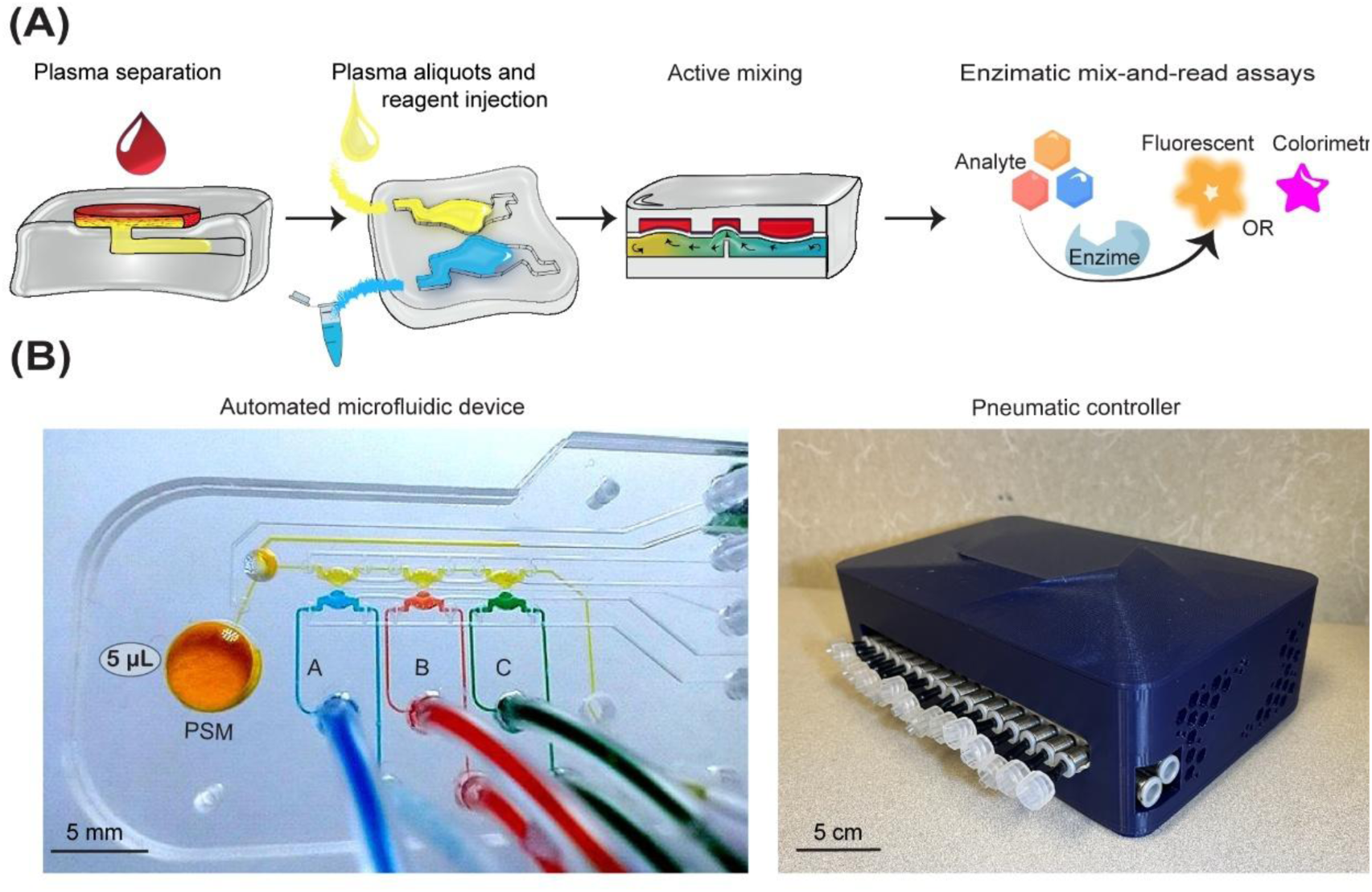
Automated thermoplastic microfluidic device. (A) Five microliters of whole blood are loaded into the device and plasma is extracted. Plasma is delivered into a bioanalysis unit where enzymatic mix-and-read assays are carried out. Detection of three analytes from the same small volume of blood is demonstrated. The device includes air microtanks to supply oxygen for enzymatic reactions. (B) The bottom panel shows pictures of the microfluidic device with colorant dyes and the pneumatic controller.

### Characterizing a microfluidic plasma separation module

Isolating plasma from whole blood is an important step in many biological assays.^25^ We wanted to develop a microfluidic plasma separation module that could be positioned upstream of the bioanalysis modules. Our ultimate goal was to isolate plasma from a finger prick of blood (5 µL) however, the resultant volume of plasma (∼1 µL) was insufficient for establishing plasma quality using commercial assays. Given that such assays typically require 10 to 20 µL of plasma, we fabricated a scaled-up version of the plasma separation module designed to extract ∼20 µL of plasma to carry out assays for bilirubin and hemoglobin. This device (shown in **Figure 2**) contained the plasma separation module but no bioanalysis modules. It had a cylindrical reservoir with a diameter of 15 mm and depth of 1.6 mm (∼100 µL volume) that served as a sample port, and an outlet for collecting plasma from the device (see **Figure 2** (**A,B**). After administering blood to the sample port, vacuum was applied at the outlet of the device and plasma began to flow into the collection microchannel, while the blood cells were retained on the PSM. The process of plasma extraction and plasma recovery from the device are outlined in **Figure S1**). Given our previous work isolating plasma in a PDMS microfluidic device, we were expecting 20% yield for plasma separation.^9^ We therefore started with ∼100 µl blood and collected ∼20 µL of plasma necessary to perform assays characterizing plasma quality.

**Figure 2.**
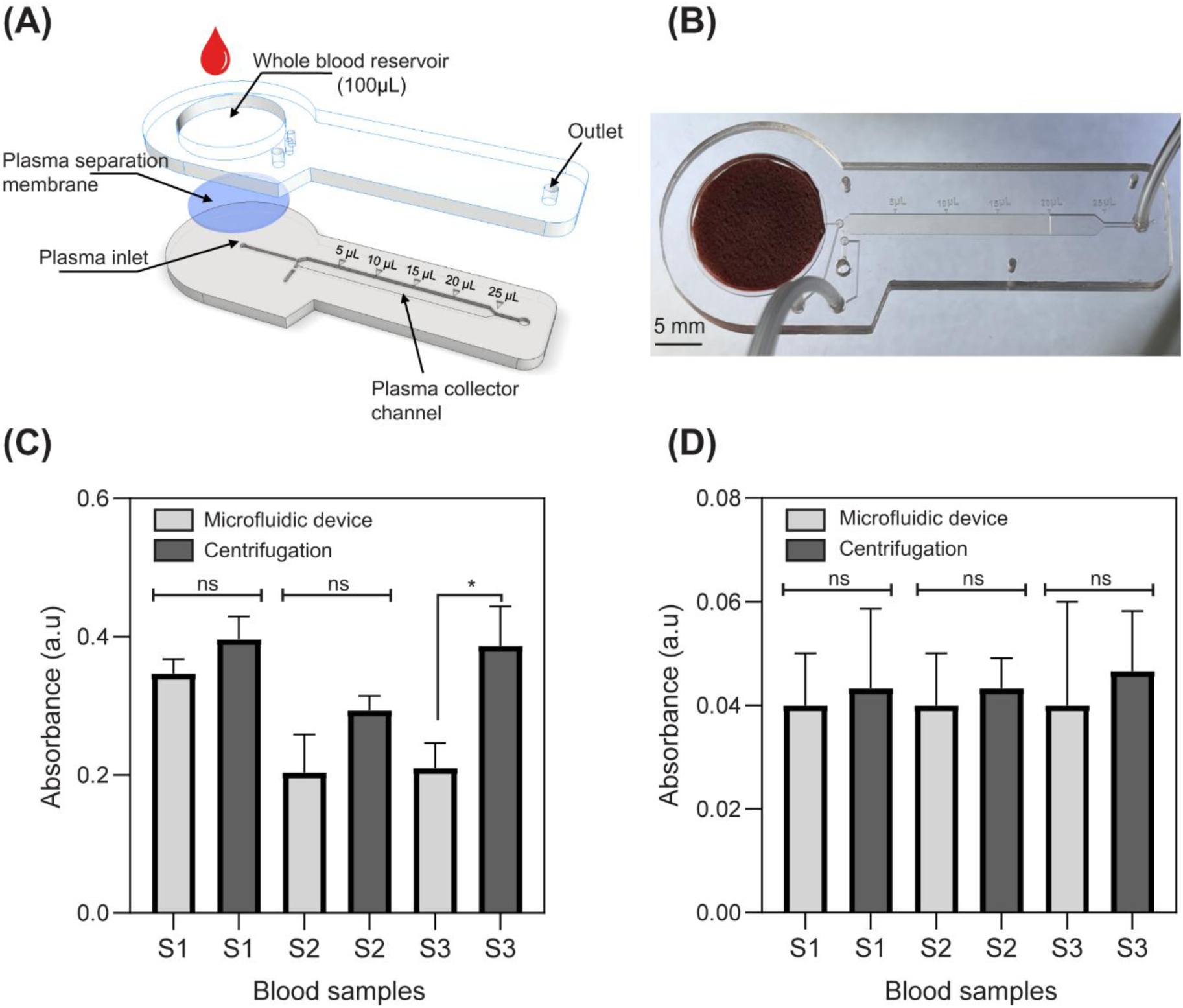
Characterizing the microfluidic plasma separator. (A) Exploded view of the components of the plasma separator device. (B) Photograph of the plasma separation device used for assessment of plasma quality. (C,D) Analysis of hemoglobin (C) and bilirubin (D) to compare quality of plasma isolated using either the microfluidic device or a conventional (centrifugation-based) method.

Hemoglobin is released during hemolysis; therefore, a lower hemoglobin level indicates a gentler separation method.^26^ Bilirubin is a byproduct of the breakdown of hemoglobin in the body and is another indicator of plasma quality.^26^ **Figure 2 (C,D)** shows hemoglobin and bilirubin analysis for three plasma samples (S1, S2, S3) extracted using either a microfluidic device or a centrifuge. Bilirubin and hemoglobin levels were similar or lower for plasma extracted by the microfluidic device compared to a centrifugation method. These data suggest that plasma extracted in our device had quality comparable to or better than the gold standard extraction method.

### Incorporating microvalves into a thermoplastic microfluidic device

On-board microvalves make it possible to automate assays while using microliter volumes of sample and reagents. However, the integration of microvalves in thermoplastic devices represents a challenge due to the high rigidity of materials (e.g., Young’s modulus of PMMA is 1800-3100 MPa). This rigidity makes it difficult to manufacture pressure-actuated valves, as these require a deformable layer to open and close effectively. Although hybrid thermoplastic valves incorporating thin flexible membranes composed of PDMS^27–30^, Viton^30^, and Fluorocur ^31^ have been developed, the process of joining layers described in these past studies was complicated and time-consuming.

Our study describes a relatively simple assembly of microvalves by joining two PMMA (acrylic) layers with PSA **(see Figure 3A**). A schematic in **Figure 3B** shows a side-view of the normally closed gate valve before and after applying negative pressure (−89.6 kPa) to the PSA. **Figure 3C** presents a top-view schematic and image of the closed gate valve. Food dye is used in the image to highlight valve architecture. We note that the gate valve was designed to withstand up to 2 kPa of flow pressure in the normally closed state, which is sufficient for most applications. We empirically converged on the design parameters including thickness of the PSA membrane (13 µm) and dimensions of the valve (0.7 mm diameter). The microchannels used in the device were 100 µm in width and depth. The microvalves were designed to minimize dead volume in the device (< 400 nL for a total of sixteen microvalves) to make it possible to analyze 1 µL of plasma.

**Figure 3.**
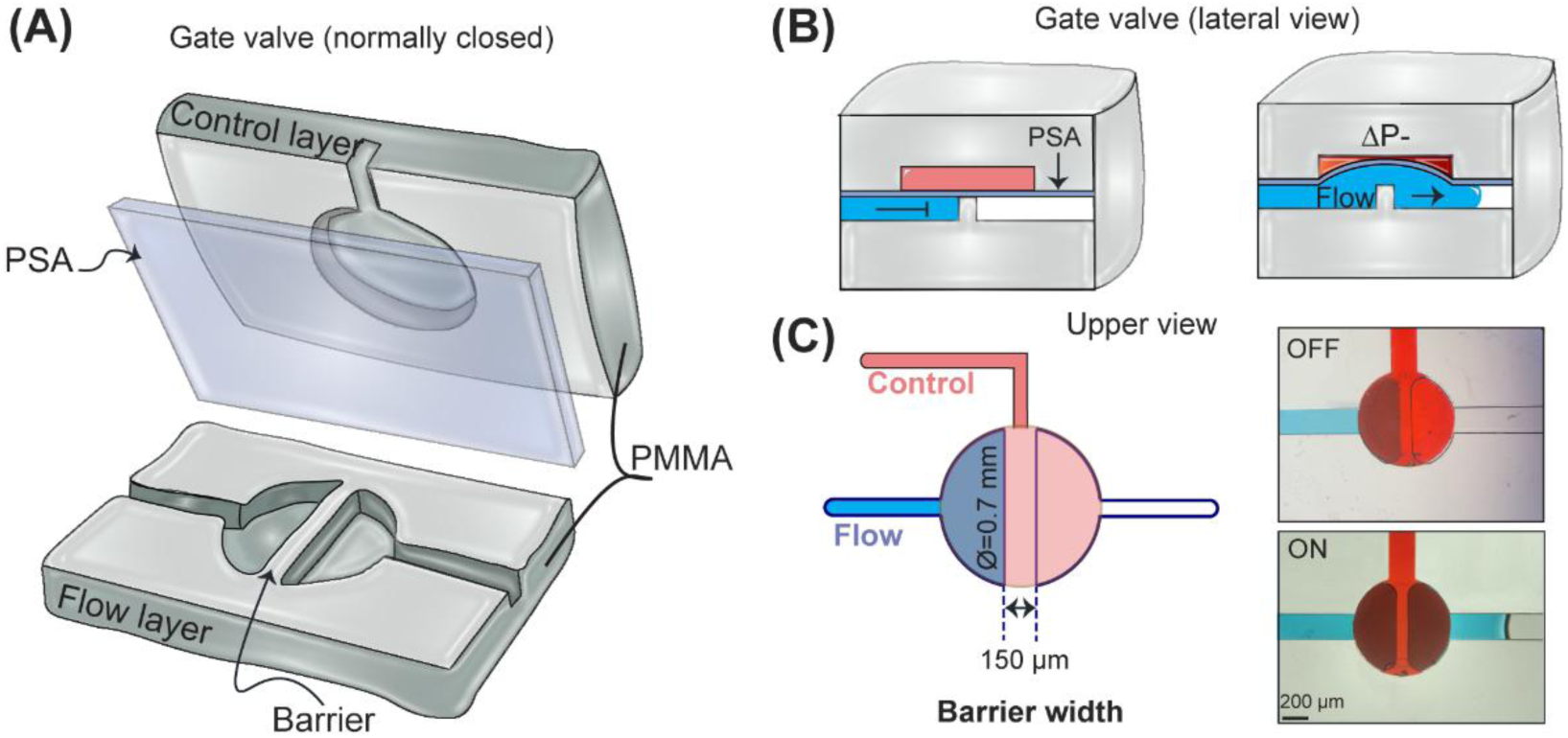
Operating microvalves. (A) 3D schematic of the normally closed gate valve. (B) Cross-sectional view of a valve in closed (off) and open (on) state. (C) Upper view of the gate valve and micrographs with blue and red food dye, indicating the flow and control layers, respectively.

### Incorporating active mixing elements into a thermoplastic microfluidic device

Mixing samples with reagents is an important step for most bioassays. Mixing in microfluidic devices is a distinct challenge because such devices typically operate under a laminar flow regime or static conditions. The challenge of mixing in microfluidic channels has been addressed by incorporating microstructures into the flow path – for example herringbone designs first described by Stroock et al.^32^ Such passive mixing elements are sufficient for a lot of microfluidic applications but are impractical for a scenario such as ours where sub-microliter volumes of samples and reagents need to be mixed rapidly. To address this challenge, we developed the concept of active mixing, where multiple PDMS microvalves operate in concert to homogenize the contents of microfluidic compartments.^9^ Active mixing allowed us to homogenize sub-microliter volumes in < 5 min regardless of molecule size. In this paper, we implemented the concept of active mixing in thermoplastic devices. As shown in **Figure 4A**, each bioanalysis unit had two valves (A and B) operated sequentially to create rapid back-and-forth movement of liquids in sample and reagent compartments. The mixing process was evaluated using food dyes. The sequence for actuating microvalves involved in active mixing is shown schematically in **Figure 4B** (left column). In this sequence we first deactivated the gate valve C by applying negative pressure (−89.6 kPa), which opened the connection between the sample and reagent compartments. This valve was kept open throughout the mixing process. Then, microvalve A was deactivated and valve B was activated by applying negative and positive pressure (−89.6 kPa, 117 kPa), respectively. The activation and deactivation of valves A and B were performed sequentially at 10 Hz for 2 min. At the end of this stage, gate valve C was closed while valves A and B were left open. At this point, image acquisition was performed. **Figure 4B** (right column) shows 4 micrographs of the bioanalysis chambers filled with yellow dye, representing plasma, and blue dye, representing reagents. As may be appreciated from this sequence of images, the contents of both chambers were homogenized after 2 min of active mixing. We note that while, as discussed in the previous section, our gate valves may operate as normally closed valves, the process of active mixing benefited from applying both negative and positive pressures.

**Figure 4.**
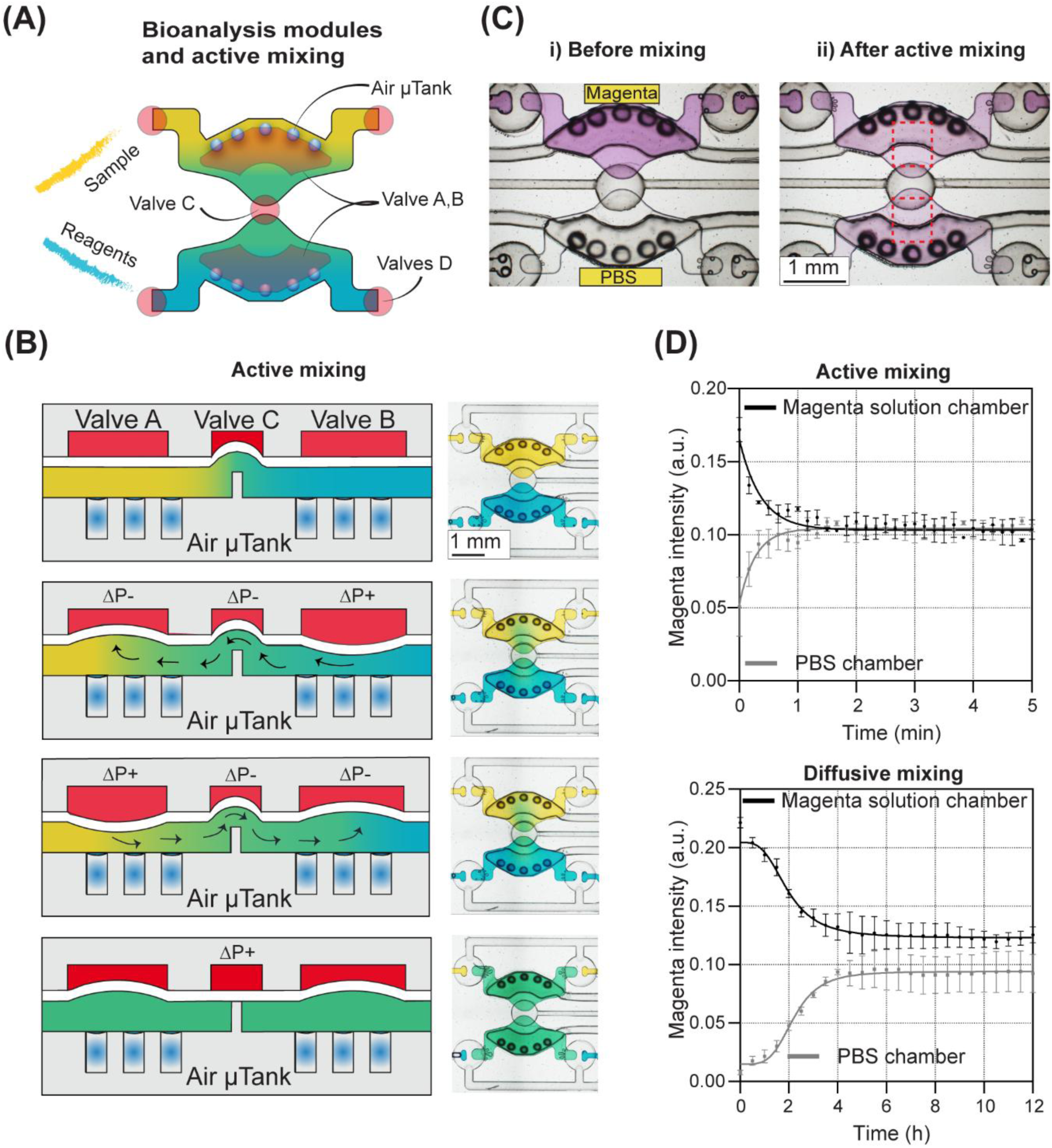
Active mixing enabled by microvalves. (A) Schematic of one bioanalysis unit showing sample and reagent chambers as well as microvalves used for active mixing (A,B,C). Flanking valves (valves D) are used to isolate each bioanalysis unit from its neighbor for independent analysis. (B) The left panel shows the cross-sectional view of the bioanalysis chambers at different stages during the active mixing. The right panel shows micrographs from a bioanalysis chamber filled with dyes at different times during mixing. The homogeneity of green color is indicative of mixing. (C) (i) Micrographs of the bioanalysis chambers loaded with assay reagents after off-chip reaction with glucose (10 mm) (upper chamber) and PBS as diluent (lower chamber). (ii) Micrographs after 2 min of active mixing. ROI used for image analysis shown in red boxes. (D) Time-lapse analysis of magenta color intensity under active mixing and diffusive mixing.

In addition to using food dye, we also evaluated the mixing process by using glucose assay reagents. We carried out glucose assay off-chip and then injected reaction products into the device. This was done to better understand how assay time is affected by mixing vs. enzymatic reaction. Optimization of the assay time involving enzymatic reaction combined with active mixing is presented in **Figure S2**. As seen from images in **Figure 4C**, homogeneity of the magenta color between sample and reagent chambers was achieved within 2 min of active mixing. Importantly, diffusion-based mixing in the absence of valve actuation took several hours to complete. The red rectangle with dashed lines in **Figure 4Cii** shows the region of interest (0.41 mm^2^) where magenta colored was analyzed. **Figure 4D** compares change in magenta color intensity in the recipient chamber (PBS chamber) over time for active and diffusion-based mixing. As highlighted by these images, maximal color intensity in the PBS chamber was achieved within 2 min for an active mixing scenario compared to 12h for a diffusion-based mixing scenario.

### Incorporating bubble traps and air micro-tanks to address challenges with gas permeability of thermoplastic devices

Air bubbles are a common challenge in microfluidic devices, primarily because air can become trapped during priming process. The confined geometry of the channels makes it difficult to displace air bubbles.^33^ Additionally, surface tension effects exacerbate the problem, as bubbles can adhere to the channel walls, particularly when the device material is hydrophobic.^33^ Because PDMS devices are gas permeable, they may be pressurized to drive bubbles out. A similar strategy for bubble removal is not possible for gas impermeable thermoplastic materials such as PMMA. **Figure 5A**, describes a hybrid bubble trap/plasma collection chamber designed to eliminate bubbles from traveling into the bioanalysis module. The bubble trap was dome shaped with a diameter of 1.6 mm and a height of 1 mm, and a volume capacity of ∼1.3 µL. Gate valves were used to control filling and emptying of the reservoir with plasma. Once sufficient volume of plasma has been collected, the inlet (channel 1) was closed off while the outlet (channel 2) was opened and plasma was delivered into the bioanalysis module (see **Figure 5A**).

**Figure 5.**
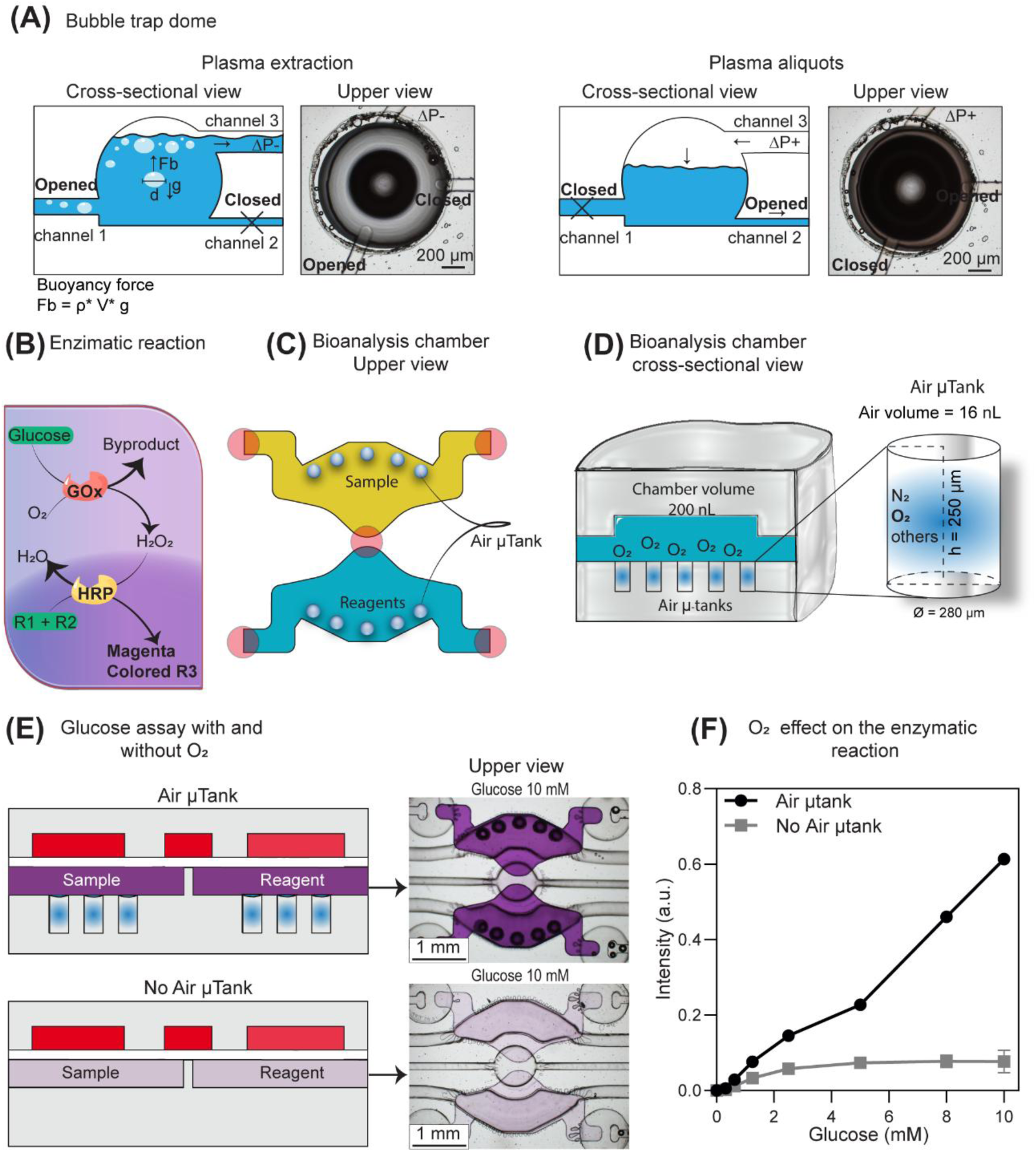
Bubble trap and air micro-tanks on chip. (A) Description of the bubble trap dome used to remove bubbles during plasma extraction and aliquoting. The microchannel 1 comes from the plasma separator, microchannel 2 is directed to the bioanalysis modules, and the microchannel 3 facilitates both the extraction and the aliquoting of the plasma. ON and OFF means valves open or closed, respectively. Fb= buoyancy force; ρ= density; g= gravitational acceleration; V= volume. (B) Enzymatic assay used for glucose detection. (C) Bioanalysis unit integrated with air micro-tanks. (C) Description of the air micro-tanks and their air storage capacities for the supply of O_2_ to the bioanalysis chambers. (D) and (E) Cross-sectional (schematic) and upper view (micrograph) of glucose assay with and without O_2_ supply. (F) The graph illustrates how the intensity of the magenta color varies in relation to the glucose concentration, under conditions with and without oxygen supplied by air micro-tanks.

A new method for bubble removal was not the only solution that needed to be developed. O_2_ is a co-factor for a large number of enzymatic reactions. For example, detection of glucose using glucose oxidase requires that O_2_ molecules are present in the vicinity of the active center of the enzyme (flavin adenine dinucleotide or FAD) to accept electrons and convert the enzyme from a reduced (inactive) to oxidized (active) state (see **Figure 5B** for description of the enzymatic reaction). When carrying out the initial glucose detection experiments in the PMMA device, we observed that linear range was limited to 2 mM glucose, much lower than in the PDMS device developed by us previously.^9^ We attributed this to differences in O_2_ permeability of the two materials and reasoned that dissolved O_2_ was insufficient to sustain enzymatic reaction in a confined volume (40 nL) of a PMMA device. To address this challenge, a PMMA microfluidic device was designed to include air micro-tanks (**see Figure 5C**) which trapped air bubbles in the floor of the sample and reagent chambers. The design of the microfluidic device, manufactured in PMMA, carefully considered both the number and volume of microtanks to provide sufficient oxygen for enzymatic reaction. **Figure 5C** shows a top view of 10 air micro-tanks integrated within each bioanalysis unit, five micro-tanks each for sample and reagent chambers. Air micro-tanks were cylindrical structures (280 µm diameter, 250 µm in length, 16 nL volume) carved in the floor of microchambers. **Figure 5D** shows a cross-sectional view of the micro-tanks in one microchamber. The total volume of air trapped in each bioanalysis unit was 0.16 µL (distributed in 10 micro-tanks). Assuming the O_2_ content of air to be 21%, the ten air micro-tanks were expected to contain ∼10 mM of O₂. Given the 1:1 stoichiometry of the enzymatic reaction, it is expected that O_2_ stored in the microtanks would be sufficient to extend the dynamic range of the glucose assay to 10 mM.

To demonstrate benefits of this design feature, we compared two versions of the automated microfluidic device, with and without air micro-tanks (see **Figure 5E**). As may be appreciated from **Figure 5F**, enzymatic reaction proceeded better, with higher sensitivity and wider dynamic range in the device with air micro-tanks.

### Operation of the automated microfluidic device and detection of glucose

After developing strategies to eliminate bubbles and supply O_2_ to the bioanalysis chambers, we proceed to detect glucose in devices. First, we carried out experiments to construct a calibration curve using known glucose concentrations (see **Figure 6 (A,B)**). The detection limits for different reaction times are described in **Figure S3.** We chose 5 min of active mixing as reaction time because it produced sufficiently low detection limit of 0.12 mM. This reaction time was used for subsequent experiments.

**Figure 6.**
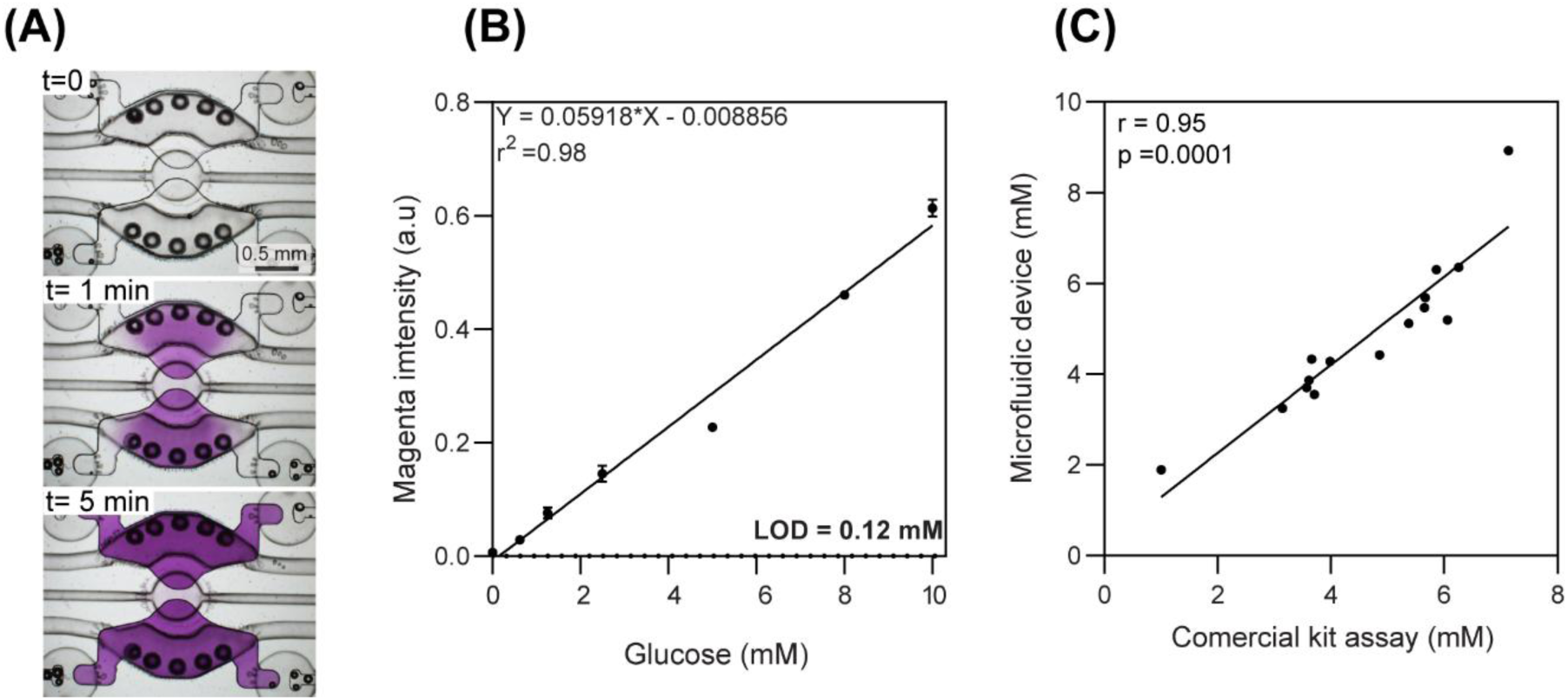
Validation of the automated microfluidic device using blood samples. (A) Time lapse of an enzymatic assay used for glucose detection. B) Calibration curve for the glucose colorimetric assay (0-10 mM) performed in the automated microfluidic device, LOD was determined by 3.3 x σ/slope, σ being the standard deviation of blank. (C) Benchmark analysis of the glucose concentration in fifteen different blood samples determined by the automated microfluidic device and a commercial glucose assay.

We validated the glucose measurements made by our automated microfluidic device against a standard method which is based on a commercial glucose kit. For this purpose, we used fifteen blood samples from healthy adult donors. For each blood sample, glucose levels were analyzed using our microfluidic device and a standard approach. For microfluidic device, 5 µL of blood was dispensed into the sample port, resulting in extraction of 1 µL of plasma. The plasma was distributed into the sample chambers of the three bioanalysis units and was mixed with glucose assay reagents to quantify glucose. For the standard assay, plasma was isolated by centrifugation and subsequently analyzed using a plate reader.

As shown in **Figure 6C**, glucose concentrations of fifteen blood samples determined in our automated microfluidic device and using a commercial assay have a close agreement with a Pearson correlation of 0.95. The coefficient of variation between measurements of the microfluidic device and standard method are less than 5%. Our results suggest that the performance of the microfluidic device is comparable to a standard method.

### Detecting three analytes from the same microliter volume of blood

As the next step, we wanted to demonstrate that multiple analytes may be detected from the same 5 µL of blood using our automated microfluidic device. To prove this concept, we added assays for lactate and H_2_O_2_ to the glucose assay described above. Measuring H_2_O_2_ is important for managing complications associated with oxidative stress whereas production of lactate may be associated with sepsis as well as hypoxia. Both biomarkers have relevance for neonatal health monitoring, ^34,35^ and both may be detected with enzymatic assays (see **Figure 7A** for description of enzymatic reactions). To showcase the broad utility of our platform, we chose to detect the former colorimetrically and later with fluorescence. Given that standard reagents are colorless, we used food dye of different colors to show the concept of multi-analyte detection. **Figure 7B** highlights our ability to load different dyes (assays) into reagent compartments while aliquoting the same amount of sample into the three sample compartments. Each sample aliquot may then be mixed with its own reagent aliquot to perform three different assays. The result of a typical analysis is shown in the micrographs of **Figure 7C**. Here, one can see colorimetric signals for lactate and glucose as well as fluorescence signal corresponding to H_2_O_2_. Calibration curves for lactate and H_2_O_2_, as well as measurements of these analytes in clinical samples are provided in **Figure S4**.

**Figure 7.**
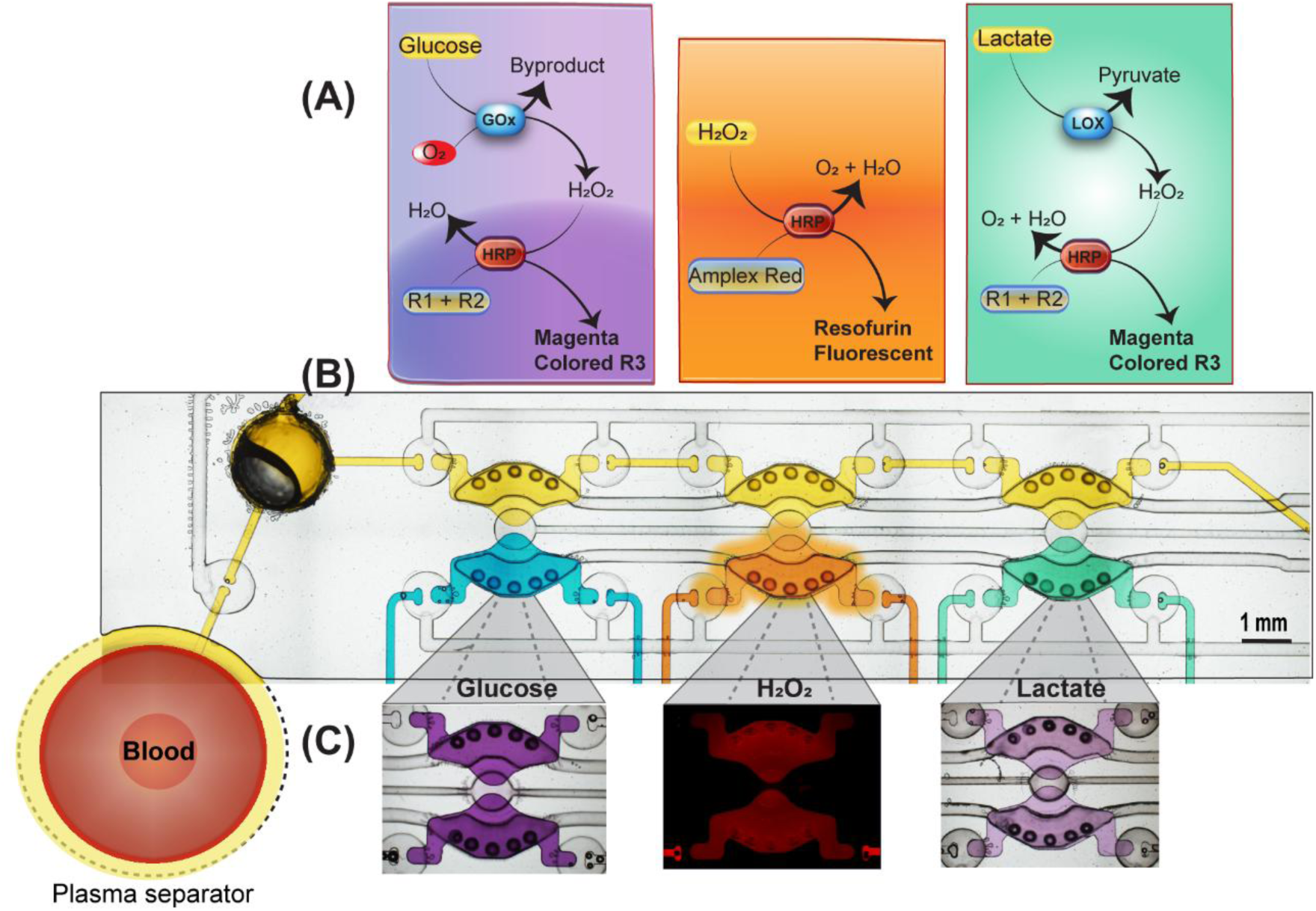
Multianalyte detection from the same small volume of blood. (A) Enzymatic assays for detection of glucose, H_2_O_2_ and lactate. Each assay is performed in its own bioanalysis unit. (B) Image of the microfluidic device microfluidic filled with yellow food dye symbolizing plasma distributed into three bioanalysis chambers. The blue, orange and green dyes symbolize the reagents for different assays to detect 3 analytes in the same volume of plasma. (C) Images of bioanalysis chambers after detection of glucose, H_2_O_2_ and lactate.

## CONCLUSIONS

This paper describes the development of an automated thermoplastic microfluidic device that performs a sophisticated workflow of plasma extraction, sample aliquoting and reagent mixing. The automation was achieved with integrated microvalves composed of PSA sandwiched between two layers of PMMA. The device also addresses challenges associated with gas impermeability of PMMA, removal of bubbles via a bubble trap and supply of O_2_ for enzymatic reactions using air micro-tanks.

The performance of the device was rigorously characterized, showing that plasma extracted in the device was comparable in quality to standard centrifugation-based isolation methods. Plasma glucose levels measured in the device were similar to values obtained with a commercial glucose detection kit. Importantly, the workflow of plasma extraction and glucose detection required only 5 µL of blood and was achieved in < 10 min. To show multiplexing capabilities of the platform, we detected lactate and H_2_O_2_, in addition to glucose. There analytes were detected using the same small volume of blood (5 µL) and short period of time (<10 min). The microfluidic device described here represents a promising technology for neonatal blood analysis and other applications where processing small volumes of blood followed by detecting multiple biomarkers is beneficial.

## ACKNOWLEDGEMENTS

This work was supported in part by the NIH grant HD100251.

## REFERENCES

(1) Camacho-Gonzalez, A.; Spearman, P. W.; Stoll, B. J. Neonatal Infectious Diseases: Evaluation of Neonatal Sepsis. Pediatr. Clin. North Am. 2013, 60 (2), 367–389. 10.1016/J.PCL.2012.12.003.

(2) Howie, S. R. C. Blood Sample Volumes in Child Health Research: Review of Safe Limits. Bull. World Health Organ. 2011, 89 (1), 46–53. 10.2471/BLT.10.080010.

(3) Lei, B. U. W.; Prow, T. W. A Review of Microsampling Techniques and Their Social Impact. Biomed. Microdevices 2019, 21 (4). 10.1007/s10544-019-0412-y.

(4) Stomnaroska-Damcevski, O.; Petkovska, E.; Jancevska, S.; Danilovski, D. Neonatal Hypoglycemia: A Continuing Debate in Definition and Management. PRILOZI 2015, 36 (3), 91–97. 10.1515/prilozi-2015-0083.

(5) Ho, H. T.; Yeung, W. K. Y.; Young, B. W. Y. Evaluation of “Point of Care” Devices in the Measurement of Low Blood Glucose in Neonatal Practice. Arch. Dis. Child. Fetal Neonatal Ed. 2004, 89 (4), 356–359. 10.1136/adc.2003.033548.

(6) Rajendran, R.; Rayman, G. Point-of-Care Blood Glucose Testing for Diabetes Care in Hospitalized Patients: An Evidence-Based Review. J. Diabetes Sci. Technol. 2014, 8 (6), 1081–1090. 10.1177/1932296814538940.

(7) Wisser, D.; Van Ackern, K.; Knoll, E.; Wisser, H.; Bertsch, T. Blood Loss from Laboratory Tests. Clin. Chem. 2003, 49 (10), 1651–1655. 10.1373/49.10.1651.

(8) Page, C.; Retter, A.; Wyncoll, D. Blood Conservation Devices in Critical Care: A Narrative Review. Ann. Intensive Care 2013, 3 (1), 1–6. 10.1186/2110-5820-3-14.

(9) Gonzalez-Suarez, A. M.; Stybayeva, G.; Carey, W. A.; Revzin, A. Automated Microfluidic System with Active Mixing Enables Rapid Analysis of Biomarkers in 5 ΜL of Whole Blood. Anal. Chem. 2022, 94 (27), 9706–9714. 10.1021/acs.analchem.2c01139.

(10) Brooks, D.; Slaughter, J. C.; Nichols, J. H.; Gregory, J. M. Reliability of Handheld Blood Glucose Monitors in Neonates: Trustworthy Arterial Readings but Capillary Results Warrant Caution for Hypoglycemia. J. Diabetes Sci. Technol. 2023. 10.1177/19322968231207861.

(11) Unger, M. A.; Chou, H. P.; Thorsen, T.; Scherer, A.; Quake, S. R. Monolithic Microfabricated Valves and Pumps by Multilayer Soft Lithography. Science. 2000, pp 113–116. 10.1126/science.288.5463.113.

(12) Song, Y.; Zhou, Y.; Zhang, K.; Fan, Z.; Zhang, F.; Wei, M. Microfluidic Programmable Strategies for Channels and Flow. Lab Chip 2024, 4483–4513. 10.1039/d4lc00423j.

(13) Busek, M.; Nøvik, S.; Aizenshtadt, A.; Amirola-Martinez, M.; Combriat, T.; Grünzner, S.; Krauss, S. Thermoplastic Elastomer (Tpe)–Poly(Methyl Methacrylate) (Pmma) Hybrid Devices for Active Pumping Pdms-Free Organ-on-a-Chip Systems. Biosensors 2021, 11 (5).

(14) Shaegh, S. A. M.; Pourmand, A.; Nabavinia, M.; Avci, H.; Tamayol, A.; Mostafalu, P.; Ghavifekr, H. B.; Aghdam, E. N.; Dokmeci, M. R.; Khademhosseini, A.; Zhang, Y. S. Rapid Prototyping of Whole-Thermoplastic Microfluidics with Built-in Microvalves Using Laser Ablation and Thermal Fusion Bonding. *Sensors Actuators*, B Chem. 2018, 255, 100–109.

(15) Weerakoon-Ratnayake, K. M.; O’Neil, C. E.; Uba, F. I.; Soper, S. A. Thermoplastic Nanofluidic Devices for Biomedical Applications. Lab Chip 2017, 17 (3), 362–381. 10.1039/c6lc01173j.

(16) Sackmann, E. K.; Fulton, A. L.; Beebe, D. J. The Present and Future Role of Microfluidics in Biomedical Research. Nature 2014, 507 (7491), 181–189. 10.1038/nature13118.

(17) Becker, H.; Gärtner, C. Polymer Microfabrication Technologies for Microfluidic Systems. Anal. Bioanal. Chem. 2008, 390 (1), 89–111. 10.1007/s00216-007-1692-2.

(18) Bhattacharjee, N.; Urrios, A.; Kang, S.; Folch, A. The Upcoming 3D-Printing Revolution in Microfluidics. Lab Chip 2016, 16 (10), 1720–1742. 10.1039/c6lc00163g.

(19) Au, A. K.; Lai, H.; Utela, B. R.; Folch, A. Microvalves and Micropumps for BioMEMS; 2011; Vol. 2. 10.3390/mi2020179.

(20) De Mello, A. Plastic Fantastic? Lab Chip 2002, 2 (2), 31–36. 10.1039/b203828p.

(21) Focke, M.; Kosse, D.; Müller, C.; Reinecke, H.; Zengerle, R.; Von Stetten, F. Lab-on-a-Foil: Microfluidics on Thin and Flexible Films. Lab Chip 2010, 10 (11), 1365–1386. 10.1039/c001195a.

(22) Guevara-Pantoja, P. E.; Jiménez-Valdés, R. J.; García-Cordero, J. L.; Caballero-Robledo, G. A. Pressure-Actuated Monolithic Acrylic Microfluidic Valves and Pumps. Lab Chip 2018, 18 (4), 662–669. 10.1039/c7lc01337j.

(23) Zhang, W.; Lin, S.; Wang, C.; Hu, J.; Li, C.; Zhuang, Z.; Zhou, Y.; Mathies, R. A.; Yang, C. J. PMMA/PDMS Valves and Pumps for Disposable Microfluidics. Lab Chip 2009, 9 (21), 3088–3094. 10.1039/b907254c.

(24) Amador-Hernandez, J. U.; Guevara-Pantoja, P. E.; Cedillo-Alcantar, D. F.; Caballero-Robledo, G. A.; Garcia-Cordero, J. L. Millifluidic Valves and Pumps Made of Tape and Plastic. Lab Chip 2023, 23 (20), 4579–4591. 10.1039/d3lc00559c.

(25) Crowley, T. A.; Pizziconi, V. Isolation of Plasma from Whole Blood Using Planar Microfilters for Lab-on-a-Chip Applications. Lab Chip 2005, 5 (9), 922–929. 10.1039/b502930a.

(26) Westerman, M.; Pizzey, A.; Hirschman, J.; Cerino, M.; Well-Weiner, Y.; Ramotar, P.; Eze, A.; Lawrie, A.; Purdy, G.; Mackie, I.; Porter, J. Plasma Hemoglobin: Potential Sources. Blood 2006, 108 (11), 3814–3814. 10.1182/BLOOD.V108.11.3814.3814.

(27) Back, J. Y.; Park, J. Y.; Ju, J. Il; Lee, T. S.; Lee, S. H. A Pneumatically Controllable Flexible and Polymeric Microfluidic Valve Fabricated via in Situ Development. J. Micromechanics Microengineering 2005, 15 (5), 1015–1020. 10.1088/0960-1317/15/5/017.

(28) Norouzi, A. R.; Nikfarjam, A.; Hajghassem, H. PDMS–PMMA Bonding Improvement Using SiO2 Intermediate Layer and Its Application in Fabricating Gas Micro Valves. Microsyst. Technol. 2018, 24 (6), 2727–2736. 10.1007/s00542-017-3676-2.

(29) Toh, A. G. G.; Wang, Z. F.; Ng, S. H. Fabrication of Embedded Microvalve on PMMA Microfluidic Devices through Surface Functionalization. In 2008 Symposium on Design, Test, Integration and Packaging of MEMS/MOEMS; IEEE, 2008; pp 267–272. 10.1109/DTIP.2008.4752998.

(30) Ogilvie, I. R. G.; Sieben, V. J.; Cortese, B.; Mowlem, M. C.; Morgan, H. Chemically Resistant Microfluidic Valves from Viton® Membranes Bonded to COC and PMMA. Lab Chip 2011, 11 (14), 2455–2459. 10.1039/c1lc20069k.

(31) Willis, P. A.; Greer, F.; Lee, M. C.; Smith, J. A.; White, V. E.; Grunthaner, F. J.; Sprague, J. J.; Rolland, J. P. Monolithic Photolithographically Patterned Fluorocur^TM^ PFPE Membrane Valves and Pumps for in Situ Planetary Exploration. Lab Chip 2008, 8 (7), 1024–1026. 10.1039/b804265a.

(32) Stroock, A. D.; Dertinger, S. K. W.; Ajdari, A.; Mezić, I.; Stone, H. A.; Whitesides, G. M. Chaotic Mixer for Microchannels. Science 2002, 295 (5555), 647–651. 10.1126/SCIENCE.1066238.

(33) Gao, Y.; Wu, M.; Lin, Y.; Xu, J. Trapping and Control of Bubbles in Various Microfluidic Applications. Lab Chip 2020, 20 (24), 4512–4527. 10.1039/d0lc00906g.

(34) Millán, I.; Piñero-Ramos, J. D.; Lara, I.; Parra-Llorca, A.; Torres-Cuevas, I.; Vento, M. Oxidative Stress in the Newborn Period: Useful Biomarkers in the Clinical Setting. Antioxidants 2018, 7 (12), 1–13. 10.3390/antiox7120193.

(35) Yilmaz, A.; Kaya, N.; Gonen, I.; Uygur, A.; Perk, Y.; Vural, M. Evaluating of Neonatal Early Onset Sepsis through Lactate and Base Excess Monitoring. Sci. Reports 2023 131 2023, 13 (1), 1–8. 10.1038/s41598-023-41776-0.

