## Supplementary material for "Integrating microfluidic automation into thermoplastic devices for analysis of small volumes of blood": SI

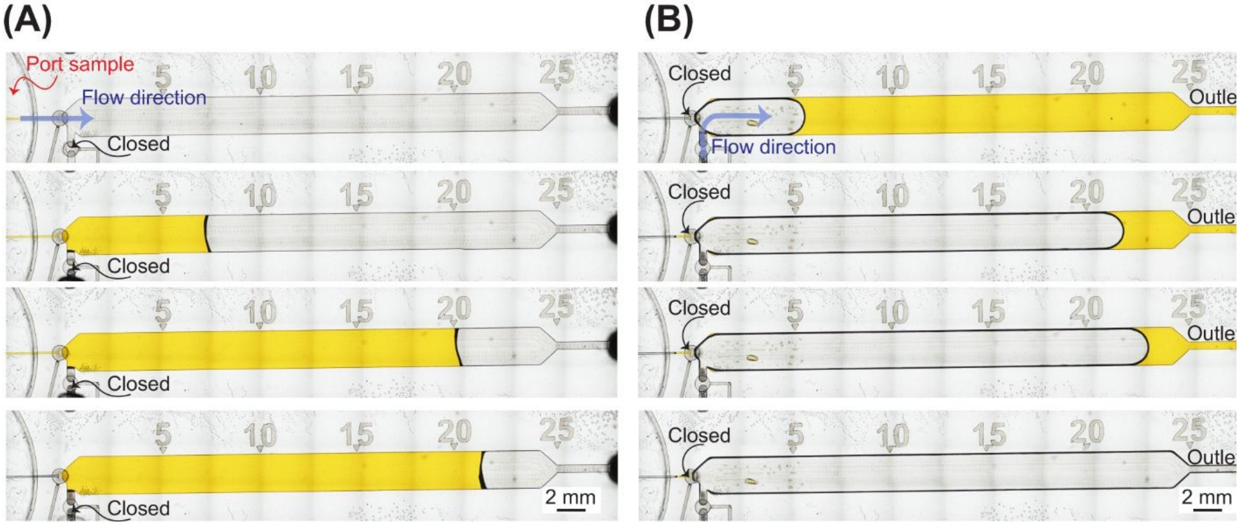

**Figure S1. Plasma separation process.** (A) Plasma extraction from whole blood. (B) Plasma recovery from the device.

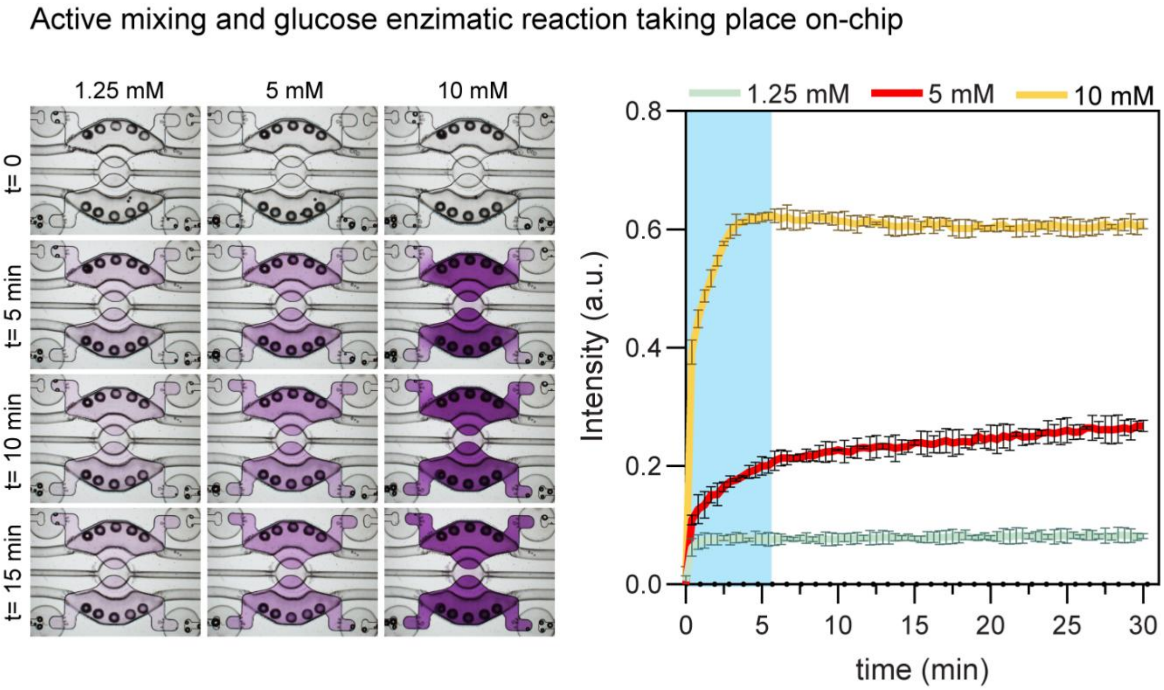

**Figure S2.** Characterization of the enzymatic oxidation of glucose over time at varying glucose concentrations.

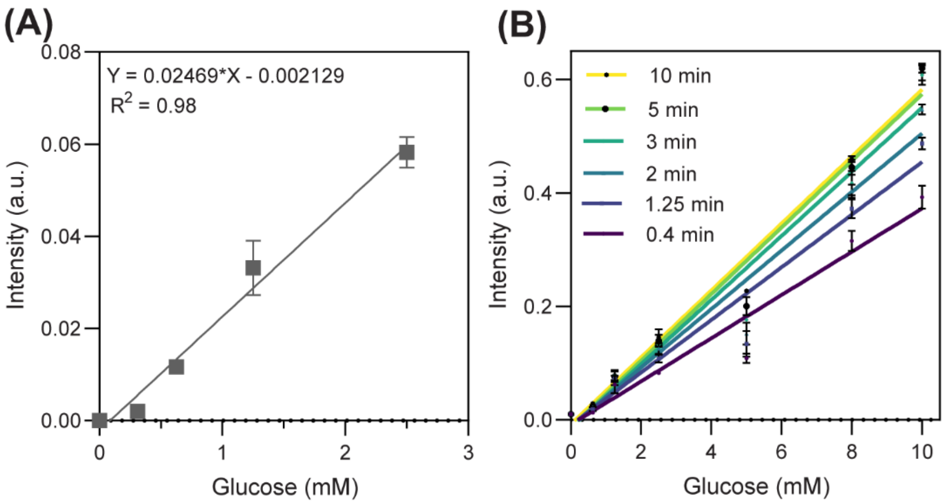


(C)

| Time (min) | LOD (Mm) |
|---|---|
| 15 | 0.12 |
| 10 | 0.12 |
| 5 | 0.12 |
| 3 | 0.13 |
| 2 | 0.14 |
| 1.25 | 0.15 |
| 0.4 | 0.19 |

**Figure S3. Linear range for glucose detection with and without air microtanks.** (A) Linear range for enzymatic glucose assay in the device without (A) and with (B) air microtanks. (C) Glucose limit of detection for different times of active mixing.

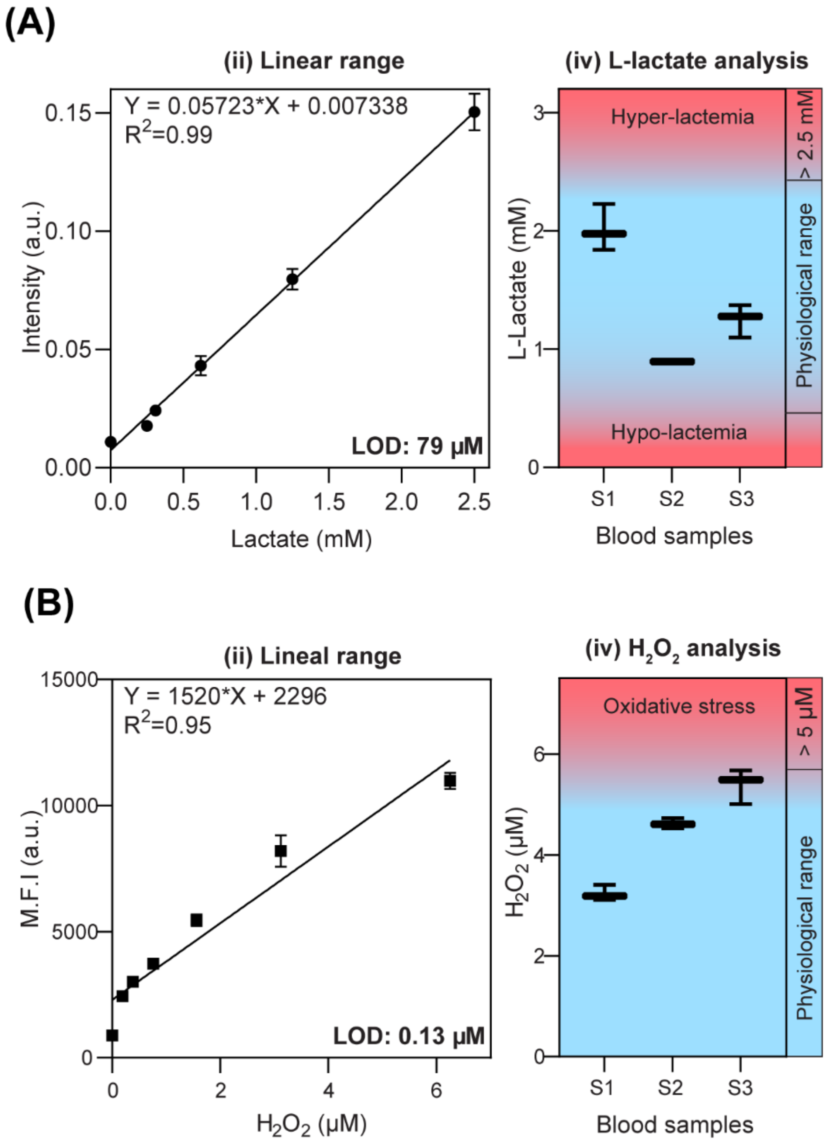


**Figure S4. Lactate and $H_2O_2$ analysis.** (A) Linear range for lactate colorimetric assay and measurements of this analyte in three clinical samples. (B) Linear range for $H_2O_2$ fluorescent assay and measurements of this analyte in three clinical samples.

**SM1. Estimation of the amount of oxygen that micro air tanks can supply to enzymatic reactions**

**Dimensions and volumes**

Microwell diameter: 0.2850 mm

Area: 0.0638 $mm^2$

Height: 0.25 mm

Air volume: 0.016 µL (air/ microwell) = $1.6*10^{-8}$ L

Air volume x 10 microwells: 0.16 µL (air) = $1.6*10^{-7}$ L

$O_2$ Volume= 0.16 (0.21) = 0.0336 µL ($O_2$) = $3.36*10^{-8}$ L

**Oxygen molarity on the 10 microwells**.

We know that the oxygen concentration in the air is 21% by volume and that 1 mole of gas occupies 24.45 L at 25°C and 1 atm.

We use the Ideal Gas Law to calculate the number of moles of air in the cylinders:

$$\mathrm{n(air)} = \frac{PV}{RT} = \frac{1(1.6 * 10^{-7})}{(0.0821)(298)} = 6.52 * 10^{-9}$$

Oxygen accounts for 21% of air, so the number of moles of oxygen is:

$n(O_2)$ =(0.21)*($6.52 * 10^{-9}$)= $1.37*10^{-9}$ mol = $1.37*10^{-9}$ mmol

**oxygen concentration**

$$CO2 = \frac{O2\ molarity}{Total\ volume} = \frac{1.37 * 10^{-9}\ \mathrm{mmol}}{1.6 * 10^{-7}\ \mathrm{L}} = 9\ mMol/L$$

In addition to the oxygen supplied by the air microtanks, the oxygen dissolved in the aqueous solution also contributes to the reaction. Since the stoichiometric ratio between glucose and oxygen is 1:1 (D-glucose + $O_2$ → D-glucono-1,5-lactone + $H_2O_2$), theoretically, the oxygen supplied by micro air tanks could detect ~ **10 mM** of glucose which ensures an adequate supply to extend the linear detection range from 2 to 10 mM.
